# Engineering Rubisco Condensation in Chloroplasts to Manipulate Plant Photosynthesis

**DOI:** 10.1101/2024.09.16.613299

**Authors:** Taiyu Chen, Marta Hojka, Philip Davey, Yaqi Sun, Fei Zhou, Tracy Lawson, Peter J. Nixon, Yongjun Lin, Lu-Ning Liu

## Abstract

Although Rubisco is the most abundant enzyme globally, it is inefficient for carbon fixation because of its low turnover rate and limited ability to distinguish CO_2_ and O_2_, especially under high O_2_ conditions. To address these limitations, phytoplankton, including cyanobacteria and algae, have evolved CO_2_-concentrating mechanisms (CCM) that involve compartmentalizing Rubisco within specific structures, such as carboxysomes in cyanobacteria or pyrenoids in algae. Engineering plant chloroplasts to establish similar structures for compartmentalizing Rubisco has attracted increasing interest for improving photosynthesis and carbon assimilation in crop plants. Here, we present a method to effectively induce the condensation of endogenous Rubisco within tobacco (*Nicotiana tabacum*) chloroplasts by genetically fusing superfolder green fluorescent protein (sfGFP) to the tobacco Rubisco large subunit (RbcL). By leveraging the intrinsic oligomerization feature of sfGFP, we successfully created pyrenoid-like Rubisco condensates that display dynamic, liquid-like properties within chloroplasts without affecting Rubisco assembly and catalytic function. The transgenic tobacco plants demonstrated comparable autotrophic growth rates and full life cycles in ambient air relative to the wild-type plants. Our study offers a promising strategy for modulating endogenous Rubisco assembly and spatial organization in plant chloroplasts via phase separation, which provides the foundation for generating synthetic organelle-like structures for carbon fixation, such as carboxysomes and pyrenoids, to optimize photosynthetic efficiency.

## Introduction

Carbohydrates are the cornerstone of life on Earth, and virtually all carbohydrates are produced directly or indirectly by ribulose-1,5-bisphosphate carboxylase/oxygenase (Rubisco) via the Calvin-Benson-Bassham (CBB) cycle (Bracher et al., 2017). As the most abundant enzyme on the planet, Rubisco fixes approximately 250 billion tons of carbon dioxide (CO_2_) per year (Field et al., 1998). Despite its crucial role in global carbon fixation, Rubisco remains a remarkably inefficient enzyme. First, Rubisco exhibits a slow turnover, catalyzing only one to ten CO_2_ fixation reactions per second (Flamholz et al., 2019). Second, Rubisco catalyzes a competing oxygenation reaction that utilizes O_2_ instead of CO_2_ (Hennacy and Jonikas, 2020a; Liu, 2022) with carboxylase and oxygenase activities dependent upon the ratio of CO_2_ and O_2_ (Andersson and Backlund, 2008). Rubisco evolved more than 3.5 billion years ago in an atmosphere that had high CO_2_ and negligible O_2_ concentrations. Thus, the poor discrimination between CO_2_ and O_2_ was not critical for Rubisco during its early stage of evolution (Berner, 2003; Nisbet et al., 2007; Tcherkez et al., 2006; Whitney et al., 2011). However, with the evolution of oxygenic photosynthesis, the concentrations of CO_2_ and O_2_ in the air have undergone significant changes, leading to an atmosphere composed of abundant O_2_ (∼ 21%) and relatively low levels of CO_2_ (∼ 0.04%) (Flamholz et al., 2019). As the O_2_ concentration increases, Rubisco exhibits a higher oxygenase activity, resulting in a reduction of approximately 30% in the net carbon fixation performed by Rubisco (Sharwood, 2017).

To compensate for the low efficiency and specificity of Rubisco and to enhance carbon fixation, photosynthetic organisms synthesize high quantities of Rubisco *in vivo* (Bainbridge et al., 1995) and have evolved various CO_2_-concentrating mechanisms (CCMs). The presence of CCMs result in different spatial organization of Rubisco in the cells, which is generally driven by liquid-liquid phase separation. In cyanobacteria and algae, Rubisco is condensed into compartmenting structures known as carboxysomes and pyrenoids, respectively, as part of their CCM (Liu, 2022; Mackinder et al., 2017). The condensation of Rubisco is mediated by multivalent interactions between Rubisco and intrinsically disordered linker proteins, including CsoS2 for α-carboxysomes, CcmM for β-carboxysomes, or Essential Pyrenoid Component 1 (EPYC1) for pyrenoids (Cai et al., 2015; Freeman Rosenzweig et al., 2017; He et al., 2020; Itakura et al., 2019; Ni et al., 2023; Oltrogge et al., 2020; Wang et al., 2019; Zang et al., 2021). In contrast, Rubisco constitutes approximately 30-50% of the soluble protein in C_3_ plant leaves and is evenly distributed in the chloroplasts, where oxygenic photosynthesis occurs (Chen et al., 2023a). Rubisco is the rate-limiting enzyme in photosynthesis, particularly in C_3_ plants that lack CCMs, including many important crops.

Given the increase in global population and climate change, there is a pressing need to explore a practical approach to increase agricultural yields (Bailey-Serres et al., 2019; Bracher et al., 2017; Croce et al., 2024; Ort et al., 2015; Whitney et al., 2011; Zhu et al., 2010). Introducing a functional CCM into C_3_ crops is considered an attractive approach for substantially improving CO_2_-fixation efficiency and plant growth (Chen et al., 2023a; Hennacy and Jonikas, 2020b; Long et al., 2015; Rae et al., 2017). An important step is to establish Rubisco condensation mediated by linker proteins in plant chloroplasts. Previous studies have reported the aggregation of cyanobacterial Rubisco mediated by CcmM35 in tobacco (Lin et al., 2014) and the condensation of plant-algal hybrid Rubisco mediated by EPYC1 into a single phase-separated compartment in the *Arabidopsis* chloroplast (Atkinson et al., 2020; Atkinson et al., 2019). However, there are still many challenges in inducing native Rubisco condensation in plant chloroplasts. Rubisco is a hexadecamer containing eight large (RbcL) and eight small (RbcS) subunits, which are encoded by the chloroplast genome and gene family in the nuclear genome, respectively (Spreitzer and Salvucci, 2002). This potentially increases the difficulty in manipulating endogenous Rubisco in plants. For example, multiple copies of *rbcS* in the plant nuclear genome need to be deleted and replaced by the *rbcS* gene of *Chlamydomonas* to generate hybrid Rubisco for condensation driven by EPYC1-RbcS interactions (Atkinson et al., 2017). Additionally, the conserved residues of native Rubisco should be modified to ensure functional interactions with non-native linker proteins (CsoS2 for α-carboxysomes and CcmM for β-carboxysomes); however, point mutations of the RbcL subunit often decrease enzymatic activity (Lin et al., 2021).

Here, we developed an approach to induce condensation of endogenous Rubisco in the chloroplasts of the model tobacco crop *Nicotiana tabacum* (*Nt*) by genetically fusing superfolder green fluorescent protein (sfGFP) to the C-terminal of tobacco RbcL through chloroplast transformation technology. The intrinsic aggregation property of sfGFP drives the formation of pyrenoid-like Rubisco condensates without affecting the assembly and enzymatic functionality of Rubisco. Rubisco condensates within chloroplasts exhibit highly dynamic liquid-like features. The transgenic tobacco plants demonstrated comparable rates of autotrophic growth to wild-type (WT) plants in full life cycles in ambient air. This study provides a strategy for plant biotechnology to conjugate plant Rubisco to form shell-less microcompartments. It lays the groundwork for developing synthetic structures that mimic natural CO_2_-fixing organelles, for example, carboxysomes and pyrenoids, which may inform plant engineering for enhancing the photosynthetic efficiency and productivity of crop plants.

## Results

### Generation of *Nt*RbcL-sfGFP transplastomic tobacco lines

As GFP is a bright fluorescent protein with a weak dimerization tendency, it has been used for oligomeric scaffolding in synthetic biology (Leibly et al., 2015). We reasoned that the intrinsic dimerization of the GFP tag that is fused to Rubisco subunits might trigger the assembly of Rubisco to a certain extent in the chloroplast. To test this, we fused sfGFP to the C-terminus of tobacco RbcL (*Nt*RbcL) using a chloroplast transformation vector (psfGFP), which is driven by *Prrn* (rRNA operon promoter) and contains *sfgfp*, the terminator *AtTpetD*, and the *aadA* gene, which encodes aminoglycoside-3’-adenylyltransferase (Figure 1a). The vector was transformed into tobacco chloroplasts using biolistic particle bombardment. After two rounds of selection and regeneration, three independent transgenic lines were obtained from the transformation, and the transgenic plants were grown autotrophically in soil and air conditions for flowering and seed maturation.

**Figure 1.**
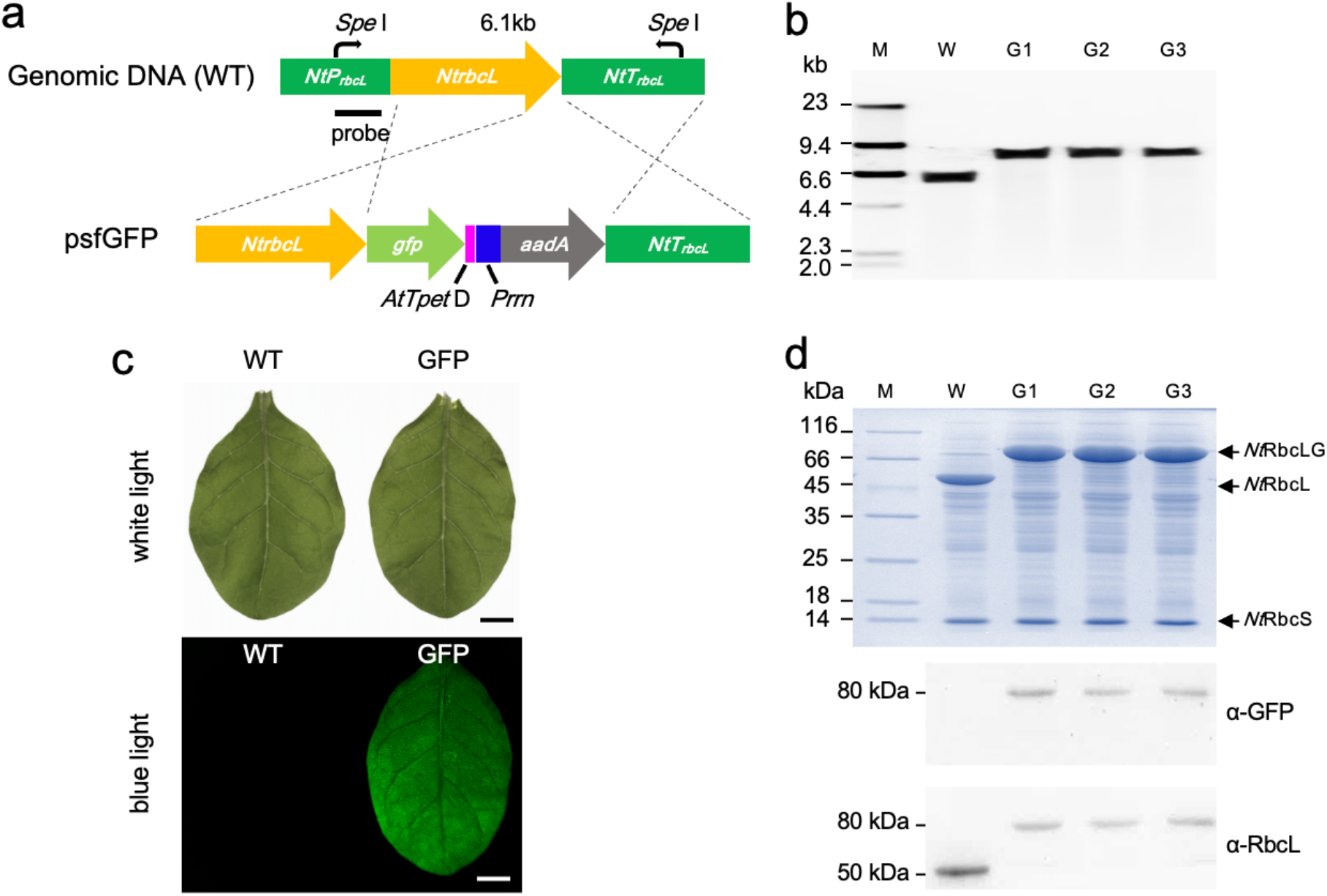
Generation of *Nt*Rubisco-sfGFP line by chloroplast transformation. **a**, Gene organization of chloroplast transformation construct (psfGFP) and the *rbcL* locus in the wild-type (WT) tobacco chloroplast genome. The stop codon in the 3’ end of *NtrbcL* was removed and fused with the coding sequence of sfGFP followed with a terminator, *AtTpet* D. The antibiotic gene (*aadA*) was driven by promoter of Prrn. IEE, Intercistronic Expression Elements; SD, Shine-Dalgarno sequence; T, Terminators. *At* indicated *Arabidopsis thaliana*. **b**, Southern blot of total genomic DNA of WT and three transgenic lines (G1 to G3). The genomic was digested by SpeI and hybridized with the probe indicated in a. The completely shifting of the fragment length polymorphism verified the successful transgene integration and homoplasmy of the three transplastomic plants generated. **c**, The leaves of transgenic plants (GFP) exhibited a robust green fluorescence signal in blue light indicating an expression of sfGFP. Scale bar: 1 cm. **d**, SDS-PAGE (top) and immunoblot analysis (bottom) of total soluble proteins confirmed the successful expression and solubility of RbcL-GFP in transgenic lines. The size of large subunit was shifted from ∼ 50 kDa (NtrbcL) to ∼78 kDa (RbcL-GFP) in transgenic line.

Southern blot analysis using the *NtrbcL* promoter as the probe showed a clear difference in fragment length between the WT and transgenic tobacco lines. (Figure 1b). The bands of the WT size (∼ 6.1 kbp) completely disappeared and were replaced with a band of ∼ 8 kbp length, indicating full replacement of WT fragments in the transgenic lines, resulting in homoplasmic transformants (Figure 1b).

### Expression of *Nt*RbcL-sfGFP in tobacco chloroplasts

To determine the expression and fusion of *Nt*RbcL-sfGFP (*Nt*RbcLG), sfGFP fluorescence was first screened under white and blue light (Figure 1c). No remarkable differences in morphology were observed between WT and transgenic leaves under white light (Figure 1c). When illuminated with blue light, the entire leaf of the transgenic plant exhibited strong green fluorescence, whereas the WT leaves showed no detectable fluorescence (Figure 1c). These observations suggest that *Nt*RbcLG was successfully expressed in transgenic plant leaves.

The fusion of sfGFP to *Nt*RbcL was confirmed by SDS-PAGE and immunoblot analysis of total soluble proteins from tobacco leaves. SDS-PAGE revealed that the size of the *Nt*RbcL protein shifted from ∼52 kDa to ∼80 kDa in transgenic lines, in agreement with the expected size of *Nt*RbcLG, and there was no detectable difference in the sizes of *Nt*RbcS between WT and transgenic lines (Figure 1d). Immunoblot analysis using α-RbcL and α-GFP antibodies revealed the absence of the WT *Nt*RbcL band in the transgenic lines, confirming the successful fusion of sfGFP to *Nt*RbcL and the full segregation of the transgenic plants (Figure 1d).

### Assembly and activity of *Nt*Rubisco-sfGFP

To investigate the assembly state of *Nt*Rubisco complexes fused with sfGFP, we performed native polyacrylamide gel electrophoresis (native-PAGE) and immunoblot analysis of total soluble proteins from tobacco to evaluate the effect of fusing GFP on the assembly of RbcL_8_S_8_. Coomassie-Blue staining and GFP fluorescence detection by native-PAGE, as well as immunoblot analysis using α-RbcL and α-GFP, showed that the molecular mass of Rubisco fused with sfGFP shifted from ∼520 kDa to ∼750 kDa, suggesting that the fusion of sfGFP to *Nt*RbcL did not affect the formation of RbcL_8_S_8_ hexadecamers (Figure 2a-2d). Interestingly, we also observed the presence of larger assemblies of [(RbcLG_8_S_8_)_n_] (Figure 2a-2d). In contrast, no large assemblies were observed for *Nt*Rubisco without sfGFP tag (Figure 2a-2c). These results suggest that sfGFP tagging could induce physical association of *Nt*Rubisco complexes within chloroplasts.

**Figure 2.**
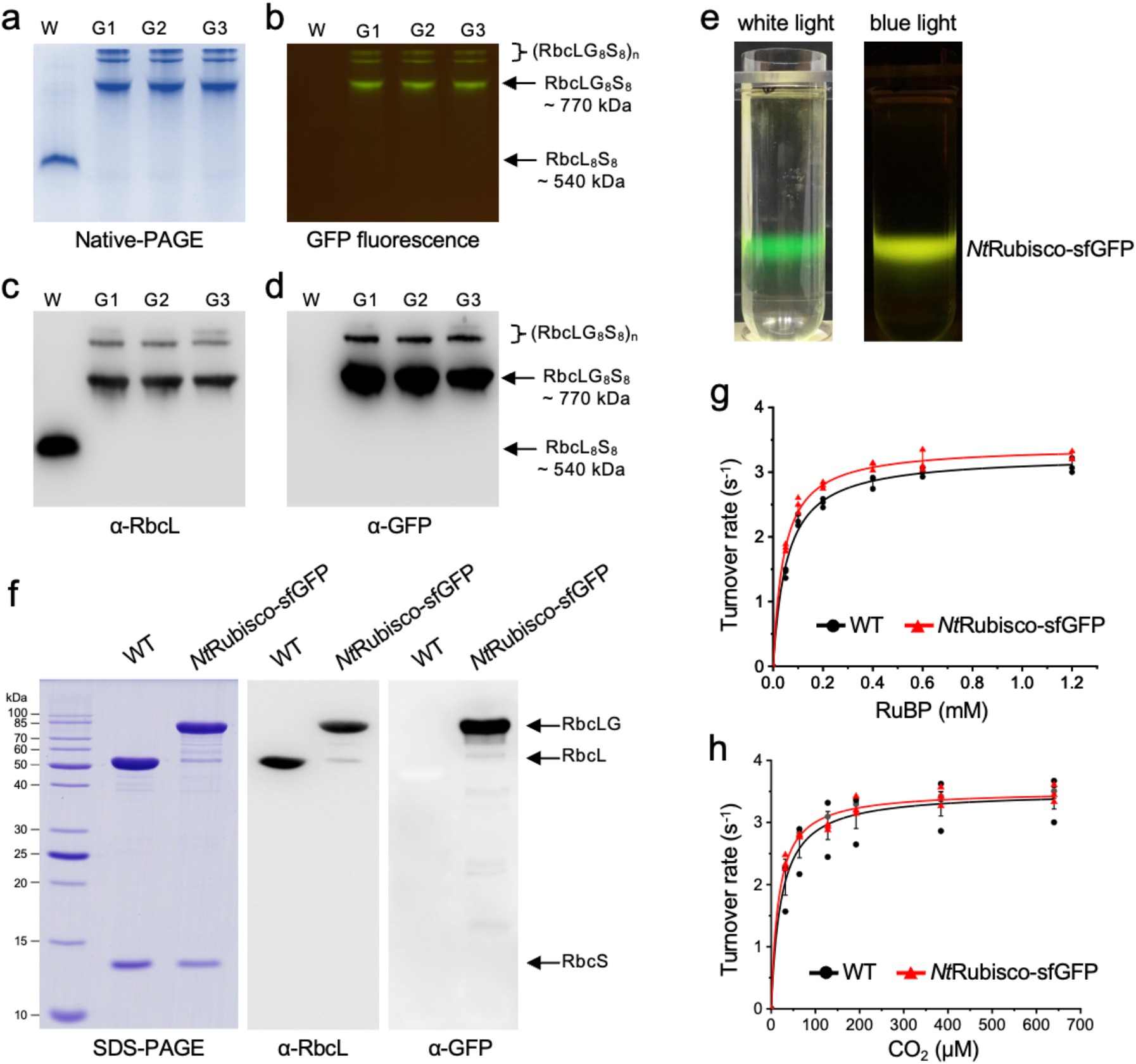
The tagging at the C terminus of NtRbcL with sfGFP did not affect the formation of L_8_S_8_ holoenzyme and enzymatic activities. **a**-**d**, Coomassie blue staining and GFP fluorescence of Native-PAGE (a and b) and immunoblot analysis using α-RbcL and α-GFP antibodies (c and d) of total soluble protein of WT and transgenic lines indicate that the expressed RbcL-GFP can assemble with Native RbcS to form Rubisco holoenzyme (∼770 kDa). The size of native Rubisco is ∼ 540 kDa and no obvious aggregated bands [(*Nt*RbcL_8_S_8_)_n_] can be detected above 540 kDa. Nevertheless, Except for the bands with ∼770 kDa, the addition of sfGFP will cause the aggregation of *Nt*Rubisco-sfGFP [(RbcLG_8_S_8_)_n_]. **e**, GFP fluorescence indicates the fraction of *Nt*Rubisco-sfGFP after the sucrose gradient centrifugation in Rubisco purification procedure. **f**, Coomassie blue staining of SDS-PAGE and immunoblot analysis using α-RbcL and α-GFP antibodies of purified Rubisco, suggesting the assembly of RbcL-GFP and NtRbcS. **g** and **h**, Rubisco activity assays as a function of different concentrations of RuBP and CO_2_ indicate that *Nt*Rubisco-sfGFP exhibit comparable enzymatic activities with that of native Rubisco (*Nt*Rubisco). The *k*_*cat*_^*C*^, *K*_*C*_, *k*_*cat*_^*R*^, and *K*_*R*_ of *Nt*Rubisco and *Nt*Rubisco-sfGFP were 3.5 ± 0.3 s^-1^ vs 3.5 ± 0.1 s^-1^, 23.4 ± 5.2 μM vs 17 ± 3.3 μM, 3.3 ± 0.1 s^-1^ vs 3.4 ± 0.1 s^-1^, and 54.8 ± 0.8 μM vs 39.7 ± 4.6 μM, respectively (*n* = 3, see Table 1). Data were fitted with Michaelis– Menten equation and are presented as mean ± SD of three independent assays.

We further purified *Nt*Rubisco-sfGFP and *Nt*Rubisco from transgenic and WT leaves, respectively, using sucrose gradient centrifugation and anion-exchange chromatography following the previously reported protocol (Carmo-Silva et al., 2011) (Figure 2e). SDS-PAGE and immunoblot analysis confirmed the presence of both RbcL and RbcS in purified *Nt*Rubisco and *Nt*Rubisco-sfGFP, as well as the fusion of sfGFP to *Nt*RbcL (Figure 2f; *Nt*RbcLG, ∼80kDa; *Nt*RbcL, ∼ 50kDa). The RbcS subunits from both the WT and transgenic lines showed similar molecular masses. In addition to *Nt*RbcLG and *Nt*RbcL, a small number of proteins with low molecular masses were detected by immunoblot analysis using an α-GFP antibody. This may be ascribed to the instability of the sfGFP tag. It is possible that during the purification process, sfGFP underwent minor non-specific degradation, resulting in the production of sfGFP and RbcL protein fragments of varying sizes (Figure 2f). Furthermore, we found that purified *Nt*Rubisco-sfGFP formed large assemblies *in vitro*, whereas free sfGFP and *Nt*Rubisco showed an even distribution in solution (Figure S1). This is consistent with our biochemical characterization results (Figure 2a-2d).

The ^14^CO_2_-fixation assays, conducted as a function of CO_2_ and ribulose 1,5-bisphosphate (RuBP) concentrations, revealed that *k*_*cat*_^*C*^ (catalytic turnover rate) and *k*_*cat*_^*R*^ (catalytic turnover rate) of *Nt*Rubisco-sfGFP were 3.5 ± 0.1 s^-1^ and 3.4 ± 0.1 s^-1^ (*n* = 3), respectively, which were similar to those of *Nt*Rubisco (3.5 ± 0.3 s^-1^ and 3.3 ± 0.1 s^-1^, *n* = 3) determined in the same assays (Table 1) and in agreement with previous findings (∼ 3.5 s^-1^) (Figures 2g and 2h) (Chen et al., 2023b; Orr et al., 2016). Additionally, *K*_*C*_ (Michaelis– Menten constant for CO_2_) of *Nt*Rubisco-sfGFP (17 ± 3.3 μM, *n* = 3) was comparable to that of *Nt*Rubisco (23.4 ± 5.2 μM, *n* = 3) (Chen et al., 2023b; Orr et al., 2016), whereas *K*_*R*_ (Michaelis–Menten constant for RuBP) of *Nt*Rubisco-sfGFP (39.7 ± 4.6 μM, *n* = 3) was slightly lower than that of *Nt*Rubisco (54.8 ± 0.8 μM, *n* = 3) (Figures 2g and 2h). Nevertheless, these results demonstrated the comparable CO_2_-fixation activities of *Nt*Rubisco-sfGFP and *Nt*Rubisco.

**Table 1.**
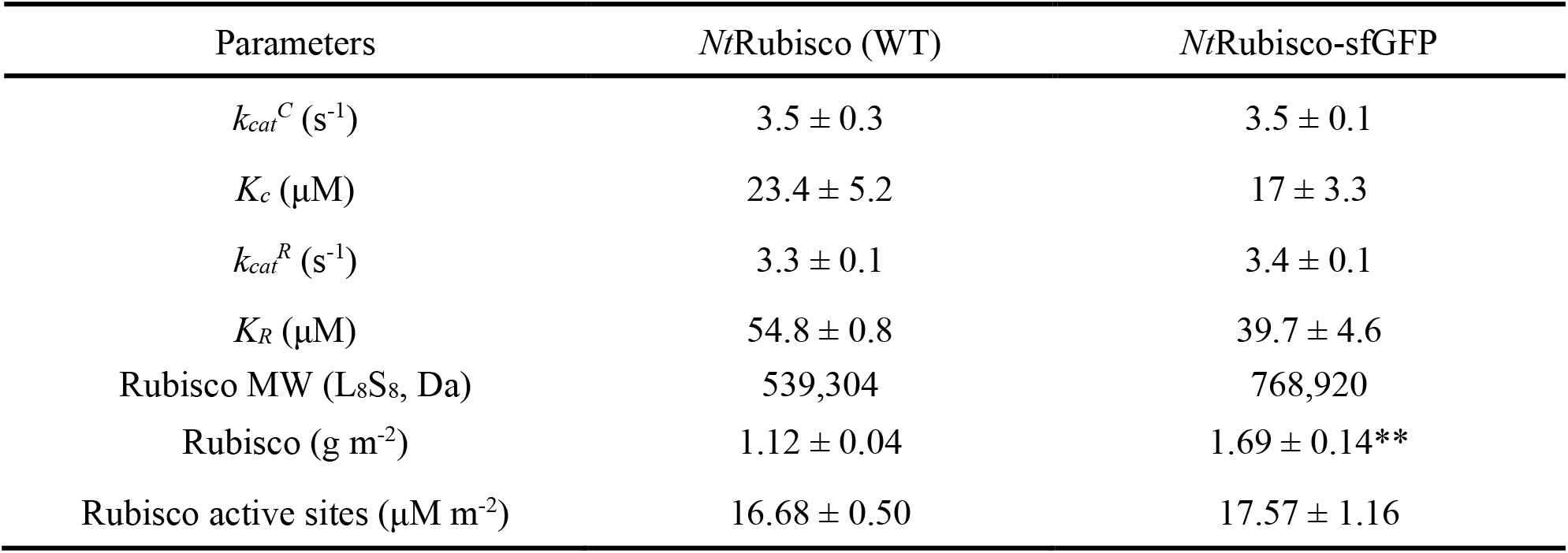
Catalytic parameters of purified Rubisco, Rubisco and chlorophyll content in *Nt*Rubisco-sfGFP and WT. Data are presented as mean ± SD (*n* = 3 independently biological replicates). **, *p* < 0.01.

### Liquid-like condensates of *Nt*Rubisco-sfGFP in tobacco chloroplasts

Rubisco complexes are evenly distributed throughout the chloroplast stroma in most plants (Atkinson et al., 2020; Chen et al., 2023a; Chen et al., 2023b). To study the location of *Nt*Rubisco-sfGFP within the chloroplast, we used confocal fluorescence microscopy to observe GFP fluorescence in the transgenic leaves. Free GFP expressed in tobacco chloroplasts exhibits an even distribution throughout the chloroplast, with its fluorescence completely colocalized with the autofluorescence of chlorophylls (Figure 3a) (Michoux et al., 2011). In contrast, the GFP fluorescence observed in the *Nt*Rubisco-sfGFP leaves exhibited an uneven distribution, with the fluorescence signal tending to accumulate in droplet-like clusters of various sizes within the chloroplasts (Figures 3b-3d). Moreover, these clusters remained stable and unaltered when exposed to the dark or 1% CO_2_ conditions (Figure S2). The maximum size of these assemblies was 3–4 µm, which is greater than that of pyrenoids (1–2 μm) but smaller than that of droplets formed by Rubisco-EPYC1 complexes *in vitro* (5– 10 μm) (Meyer et al., 2017; Wunder et al., 2018). Moreover, thin-section transmission electron microscopy (TEM) confirmed that the *Nt*Rubisco-sfGFP proteins formed a large condensate in each chloroplast (Figure S3), which is reminiscent of the dense body formed by cyanobacterial Rubisco-CCM35 complexes in tobacco chloroplasts (Lin et al., 2014) and *in vitro* observations (Figure S1).

**Figure 3.**
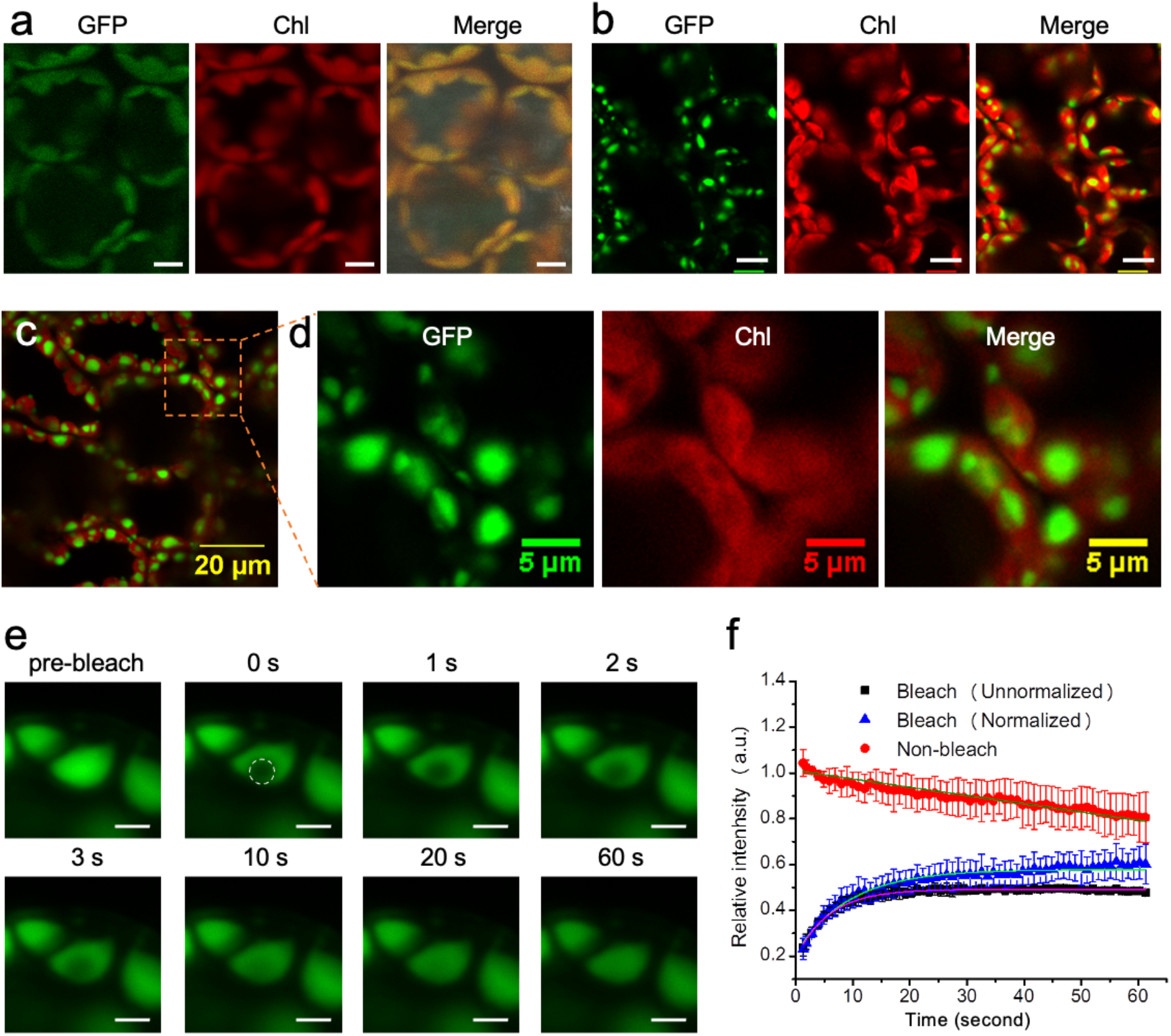
Highly dynamic mobility of Rubisco condensation directed by the dimerization of sfGFP in tobacco chloroplast. **a**, Free sfGFP exhibited an even distribution profile in tobacco chloroplast. **b**, GFP signal displayed an obvious aggregation in tobacco chloroplast. Scale bar: 10 µm. **c**, GFP signal (green) showed obvious spots rather than full co-localization with auto-fluorescence of chlorophyll in chloroplast. **d**, Higher magnification view of the area as indicated in c. Left, GFP signal; middel, auto-fluorescence of chlorophylls; right, merge image. **e**, Representative fluorescence images before/after bleaching, tracking at various time points. The rapid recovering (in 10 seconds) of the fluorescence after bleaching indicated a highly dynamic mobility of the condensation of *Nt*Rubisco-sfGFP. Scale bar: 2 µm. **f**, Representative curves of fluorescence recovery of bleached regions of the condensation of *Nt*Rubisco-sfGFP. The y-axis indicates the fluorescence intensity relative to the fluorescence intensity of the same region before bleaching. The non-bleach curve represents the fluorescence intensity of un-bleaching region. Data are presented as mean ± SD from three independent bleaching regions.

Furthermore, fluorescence recovery after photobleaching (FRAP) was exploited to evaluate the organizational status and dynamics of the condensate structures in tobacco chloroplasts (Figure 3e). The bleached fluorescence signal quickly recovered within 10 s after photobleaching, and the fluorescence reached a plateau after 40 s (Figure 3f), comparable to that of the EPYC1-induced condensates of the plant-algal hybrid Rubisco (Atkinson et al., 2020). These results revealed that the Rubisco condensates exhibited liquid-like internal mixing behavior, suggesting that the sfGFP tag condenses Rubisco complexes into a liquid matrix in the stroma of tobacco chloroplasts.

### Growth phenotypes and gas exchange assays of the *Nt*Rubisco-sfGFP transformants

To examine plant growth performance, *Nt*Rubisco-sfGFP transgenic lines were planted and grown with WT tobacco in Murashige and Skoog (MS) medium and soil. In MS medium, the *Nt*Rubisco-sfGFP plants exhibited a larger cotyledon than the WT plants during the early growth stages (Figure S4). When the plants were grown in soil, seeds of the *Nt*Rubisco-sfGFP plants germinated, and the *Nt*Rubisco-sfGFP plants demonstrated autotrophic growth and full life cycles in ambient air. There was no notable difference in growth between *Nt*Rubisco-sfGFP and WT tobacco in terms of leaf number, plant height, and dry weight (Figure 4a-4d). Moreover, there was no distinguishable difference in Rubisco content (measured by Rubisco active sites) (Table 1, Figure S5). The absolute protein mass of Rubisco was remarkably higher in the *Nt*Rubisco-sfGFP lines than in *Nt*Rubisco (Table 1), partially due to the greater molecular mass of *Nt*Rubisco-sfGFP than that of *Nt*Rubisco. Gas exchange measurements showed similar net CO_2_-assmilation rates of the *Nt*Rubisco-sfGFP lines compared to WT plants as a function of intercellular CO_2_ concentration (Ci) and light density (Figure 4e and 4f). These results indicate that the catalytic activities of *Nt*Rubisco-sfGFP in transgenic chloroplasts could support plant growth and autotrophic photosynthesis comparable to the WT tobacco plants.

**Figure 4.**
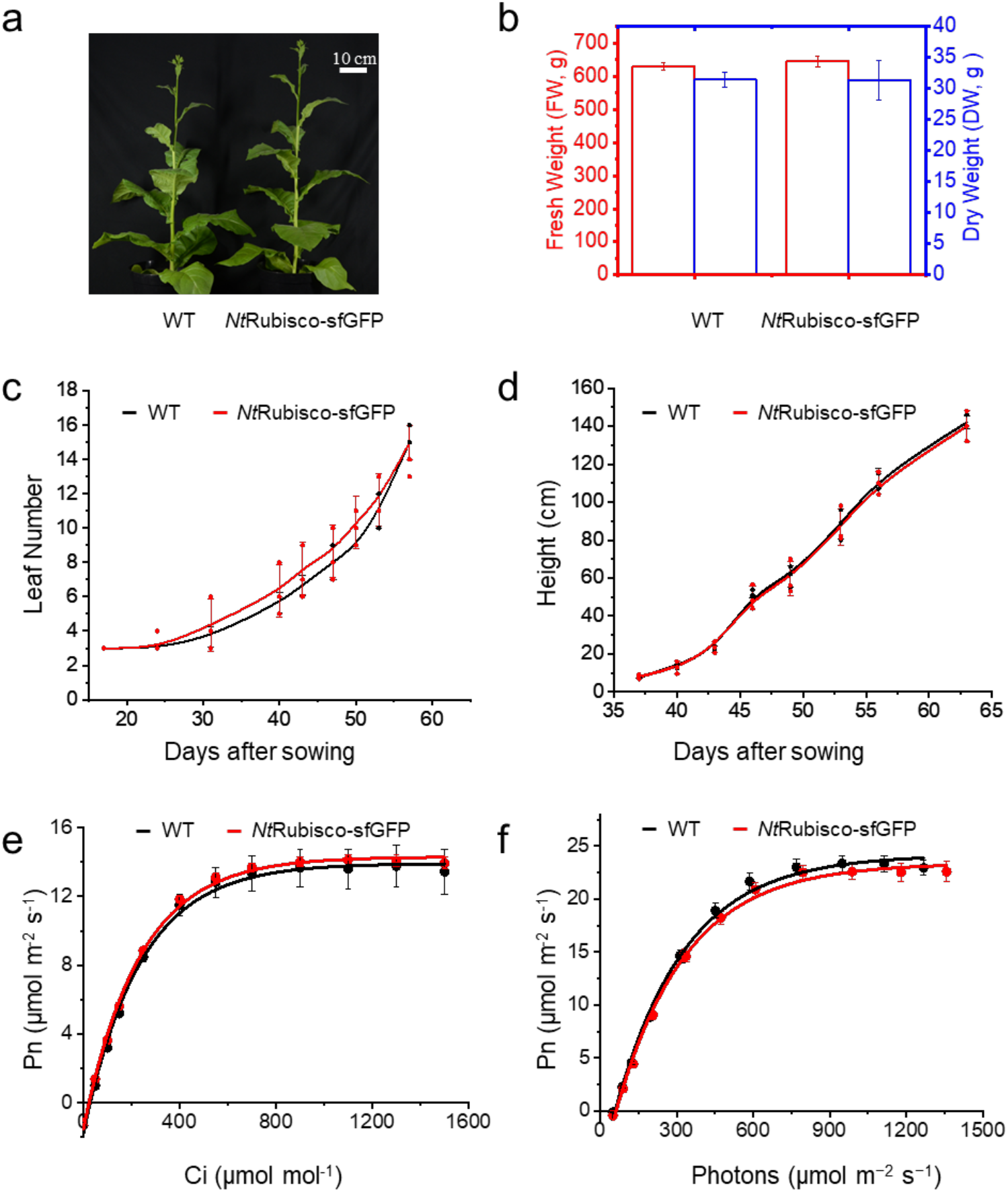
Condensate Rubisco supports autotrophic growth of tobacco plants in ambient air. **a**, Phenotypes of the *Nt*Rubisco-sfGFP and WT plants grown at flowering stage (55 days after sowing) in air condition. **b**, Fresh and dry weight of the plants showed comparable biomass of the *Nt*Rubisco-sfGFP with WT. **c** and **d**, Plant growth rate indicated as leaf number and height after sowing. **e** and **f**, Leaf gas exchange analysis of net CO_2_ photosynthesis rate (Pn) of *Nt*Rubisco-sfGFP and WT at different intercellular CO_2_ concentrations (Ci) and light densities at 25°C. Data are presented as mean ± SD (n = 3).

## Discussion

In this study, we modified the endogenous plant Rubisco in tobacco chloroplasts through genetic fusion of sfGFP to native *Nt*RbcL. As a fluorescent tag, sfGFP has been shown to facilitate protein folding and minimize protein aggregation (Scott et al., 2018). Our results showed that sfGFP tagging did not affect the assembly of tobacco Rubisco hexadecamers and their enzymatic activities within chloroplasts. Intriguingly, the self-oligomerization of sfGFP successfully induced phase separation and directed the condensation of Rubisco to form large liquid-like assemblies in the stroma of tobacco chloroplasts. The obtained transgenic plants showed autotrophic growth and a full life cycle in ambient air, comparable to the WT plants. The detailed mechanisms of *Nt*RbcL-sfGFP condensation in chloroplasts and modulation of its formation and function require further investigation.

Rubisco is a rate-limiting enzyme in photosynthesis, particularly in C_3_ crop plants that lack CCMs. Improving Rubisco catalytic performance and introducing functional CCM systems hold great promise for enhancing plant photosynthesis and productivity. However, the assembly of plant Rubisco involves multiple chaperone proteins, representing a significant challenge for Rubisco engineering (Aigner et al., 2017; Spreitzer and Salvucci, 2002). In cyanobacterial and algal CCMs, Rubisco condensation is fundamental for the construction, maintenance, and CO_2_-fixing functionality of proteinaceous organelle-like structures (carboxysomes and pyrenoids). Recent studies have shown the possibility of generating condensates of plant-algal hybrid Rubisco using pyrenoid-specific linker protein EYPC1 in *Arabidopsis* chloroplasts to support plant growth (Atkinson et al., 2020; Atkinson et al., 2019). In contrast, our study showed a new approach to establish the condensation of endogenous Rubisco complexes in plant chloroplasts, which could serve as individual physical matrices for generating advanced organelle-like structures for carbon fixation, such as carboxysomes and pyrenoids. This study may provide important information for future efforts to engineer functional CCM modules in crop plants, with the intent of improving crop photosynthesis and yield. It may also inform the bioinspired design and manipulation of protein assemblies in synthetic biology for various biotechnological and biomedical applications.

## Materials and methods

### Plant material and growth condition

The wild-type tobacco crop *Nicotiana tabacum* and transgenic lines were planted in greenhouse under normal treatment conditions. The sterile plants were germinated on the rooting medium (1/2 Murashige and Skoog medium with 3% sucrose) and were grown in an incubator (12h 25°C, day/12h 20°C night, 200 μmol photons m^-2^ s^-1^). After sterilization with ethanol, the offspring of transgenic and wild-type seeds were germinated on rooting medium with 500 mg spectinomycin for seed testing.

For growth tracking, plants were germinated and grown in a 5 L pot containing Levington Advance Seed & Modular F2S Compost and perlite (3:1, v:v). Plants were cultured in a greenhouse located at the bioscience building at the University of Liverpool, United Kingdom. Leaf number and plant height were recorded and measured during the entire life cycle to compare the growth performance of *Nt*Rubisco-sfGFP with that of wild-type (WT) plants. The fresh weight of plants was measured at the flowering stage, and the plants were then completely dried at 65°C for one week for dry weight measurement. For growth comparison, the seeds of *Nt*Rubisco-sfGFP and WT were sterilized and germinated in rooting medium, and similar growth stage seeds were selected for further growth on the plate. Images were taken at 20 days using Nikon D7000, and the sizes of leaves were further analyzed using Image J software.

### Vector construction

The chloroplast transformation vector was designed as shown in Figure 1a. In detail, nearly 850 bp of the 3’ end of *rbc L* after removing the stop codon, the cds of sfGFP (Superfolder Green Fluorescent Protein) (Pedelacq et al., 2006), and terminator *(AtTpet* D) were amplified from tobacco genomic DNA, plasmid, and Arabidopsis genomic DNA, respectively. The amplicons were then assembled to form the *rbcLgfp* cassette using Golden Gate Assembly. The selection gene (aadA) and terminator of *rbc L* were amplified from pTPTR (plasmid for Tobacco Plastid Transformation of RbcL) (Chen et al., 2023b) and built with *rbcLgfp* into pEASY^®^-Blunt Zero (Transgenbiotech, Beijing, China) using Gibson assembly (E5510S, NEB, www.neb.uk.com) to generate the final transformation vector.

### Chloroplast transformation

After coating with gold particles, the plasmid was delivered into tobacco chloroplasts following the bombardment method (Chen et al., 2023a; Chen et al., 2023b; Zhou et al., 2007). The transformed leaves were cut into 5 mm x 5 mm and selected on RMOP medium (Murashige and Skoog (MS) medium with 3% (w/v) sucrose, 500 mg L^-1^ spectinomycin, 1 mg L^-1^ 6-benzylaminopurine, 0.1 mg L^-1^ naphthaleneacetic acid, 1 mg L^-1^ thiamine-HCl, 100 mg L^-1^ Myo-inositol, 0.3% (w/v) phytagel, pH 5.8) (Svab et al., 1990). The regenerated shoots were cut and transferred to a rooting medium (1/2 MS medium with 3% (w/v) sucrose and 500 mg L^-1^ spectinomycin). Well-rooted plants were transferred into soil pots for growth, flowering, and seeding.

### Genomic DNA extraction and Southern blotting

Genomic DNA was extracted by the cetyltrimethylammonium bromide (CTAB) method as described by the reference (Chen et al., 2023a; Chen et al., 2017), and nearly 3 μg of genomic DNA was digested with *Spe* I (NEB) for Southern blotting. After separation on 0.8% agarose gel, the DNA was transferred to membranes (Amersham, http://www.amershambiosciences.com/) using the capillary method (Southern, 1975). The *rbcL* promoter region, as indicated in Figure 1, was selected as the probe for hybridization. Probe labelling, hybridization, probe washing, and immunodetection were conducted according to the manufacturer’s instructions (https://www.roche.com/). Finally, the signal was detected using an ImageQuant™ LAS 4000 system (GE Healthcare, United States).

### Confocal microscopy and Fluorescence recovery after photobleaching (FRAP)

Purified *Nt*Rubisco was labeled with Alexa Fluor 647 dye according to the previous study (Zang et al., 2021) and imaged using a Zeiss LSM780 confocal microscope with excitation and emission at 650 nm and 671 nm, respectively. The leaf samples were detached from the plants after 2 h of light, dark adaption, or 1% CO_2_ treatment, and then attached to a slide soaked with a drop of water. The samples were covered with a cover glass and imaged using a Zeiss LSM780 confocal microscope with excitation at 488 nm and 650 nm. GFP and chlorophyll fluorescence signals were measured at 500–520 nm and 660-700 nm, respectively. The GFP signal in the center of the spot was bleached with 100% laser, and images were taken every 250 ms for 60 s to track the fluorescence recovery. The fluorescence profiles were obtained and analyzed using ImageJ software, and total fluorescence was used to normalize the fluorescence distributions after bleaching.

### Rubisco purification, protein gel analysis, immunoblotting analysis, and carbon-fixation assays

Leaf discs (2 cm^2^) were quickly homogenized in extraction buffer [50 mM EPPS, 20 mM NaHCO_3_, 10 mM MgCl_2_, 1% (w/v) polyvinylpolypyrrolidone (PVPP), 5 mM dithiothreitol (DTT), and 1% protein inhibitor, pH=8.0]. Following homogenization, the mixture was centrifuged at 12,000 g for 5 min. The supernatants were used for Rubisco quantification using immunoblot analysis. *Nt*Rubisco-sfGFP and WT Rubisco were purified using the previously reported ammonium sulfate method (Carmo-Silva et al., 2011). After quantification by the Bradford method (Bradford, 1976), the protein samples were mixed with 4× SDS Sample Buffer and boiled at 100°C for 10 min for denaturing, analyzed on 15% SDS-PAGE according to the standard protocol, and then stained with Coomassie Brilliant Blue G-250 or transferred to membranes for immunoblotting analysis. Rabbit polyclonal α-rbcL (Agrisera, AS03037, dilution 1:10,000), α-GFP (Invitrogen, 33-2600, dilution 1:10,000), goat α-rabbit Immunoglobulin G (HandL), horseradish peroxidase-conjugated (Agrisera AS101461, dilution 1:10,000), and goat α-mouse secondary antibody, horseradish peroxidase-conjugated (Promega, W4021, dilution 1:10,000) were used for immunoblotting following the procedure described previously (Huang et al., 2020; Sun et al., 2016). For native-PAGE gel analysis, the samples were mixed with 4× Native Sample Buffer and separated on 7% (w/v) native-PAGE gels at 50 volts overnight. After incubation in SDS transfer buffer (0.1% (w/v) SDS, 25 mM Tris, 192 mM glycine, 20% (v/v) methanol, pH 8.3) for 1 h, protein transfer and immunoblotting analysis of native-PAGE gels were performed using the same protocol as for SDS-PAGE gels. The signal was detected using the ImageQuant™ LAS 4000 system (GE Healthcare, United States) or Tanon 5200 (Tanon, China).

Rubisco CO_2_-fixing activity was determined using the ^14^C method (Sun et al., 2016; Sun et al., 2019). Rubisco active sites were quantified by titration assay with different concentrations of carboxyarabinitol-1,5-bisphosphate (CABP) (Chen et al., 2023b). The radioactivity was measured using a scintillation counter (Tri-Carb; Perkin-Elmer, US). The CO_2_-fixing efficiency was calculated using raw data and normalized to Rubisco active sites. The Michaelis-Menten plot was fitted using Origin (www.origin.com), and all results are presented as mean ± standard deviation (SD).

### Measurement of chlorophyll content and plant weights

Leaf samples (2 cm^2^) were punched from the plants and weighed prior to chlorophyll extraction. Chlorophylls were completely extracted in 2 mL chlorophyll extraction buffer [ethanol, acetone, and water (4.5:4.5:1, v:v:v)] in the dark at 4°C overnight. The chlorophyll content was measured by the spectrophotometric method using NanoDrop Ds-11 (DeNovix, USA), and was calculated using the equations of Lichtenhaler and Wellburn (Lichtenthaler and Wellburn, 1983). The fresh shoots of plants were weighed and then dried at 60°C for one week to collect the dry weight data.

### Leaf Assimilation rates and Gas exchange

Net photosynthesis rates of fully light-adapted plants (Pn, μmolm^−2^ s^−1^) at 25°C over different concentrations of CO_2_ and photons were examined on the same leaves during gas-exchange experiments using a portable flow-through LI-6800 gas-exchange system (LI-COR, Nebraska, USA). Gas exchange data were modelled and calculated according to previous studies (Farquhar et al., 1980; von Caemmerer, 2000).

## Acknowledgments

We thank the Liverpool Biomedical Electron Microscopy Unit (Greg Dykes, Alison Becket) and the Centre for Cell Imaging for technical assistance and provision for microscopic imaging. We thank Dr. Douglas Orr and Professor Martin Parry for kindly sharing CABP as a gift. This work was supported by the National Key R&D Program of China (2023YFA0914600, 2021YFA0909600), the National Natural Science Foundation of China (32070109), the National Program of Transgenic Variety Development of China (2016ZX08001-001), the Leverhulme Trust (RPG-2021-286 to L.-N.L.), the Royal Society (URF\R\180030 to L.-N.L.), the Biotechnology and Biological Sciences Research Council Grant (BB/Y01135X/1, BB/Y008308/1, BB/V009729/1, BB/M024202/1 to L.-N.L., BB/N016807/1 to P.J.N), and the International Postdoctoral Exchange Fellowship Program from China Postdoctoral Science Foundation (20180079 to T.C.).

## Author contributions

T.C., P.N., Y.L., and L.-N.L. designed the research; T.C., M.H., D.P., S.Y. F.Z., and G.F.D. performed the research; T.C., T.L., P.N., Y.L., and L.-N.L. analyzed the data; T.C. and L.-N.L. wrote the manuscript with contributions from all other authors.

