## Supplementary Information for "Engineering Rubisco Condensation in Chloroplasts to Manipulate Plant Photosynthesis"

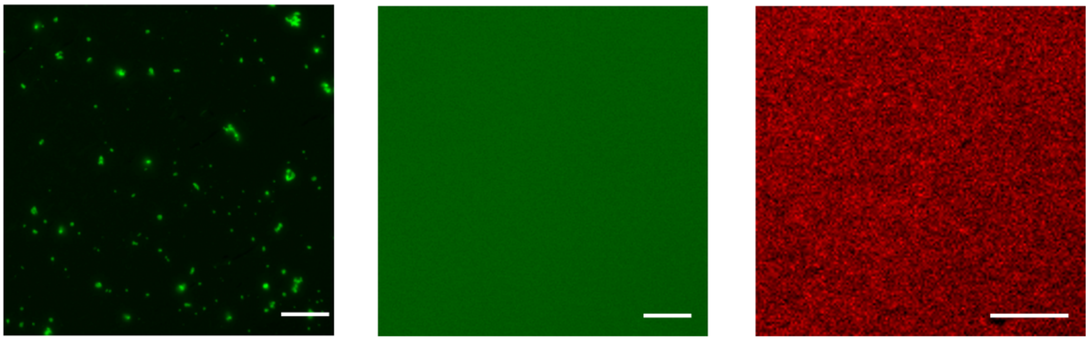


**Figure S1. *Nt*Rubisco-sfGFP purified from transgenic tobacco leaves showed obvious particle *in vitro* (Left); however, purified GFP (Middle) and *Nt*Rubisco (stained with Alexa Fluor® 647, Right) showed an even distribution *in vitro*.** Scale bar: 10 μm

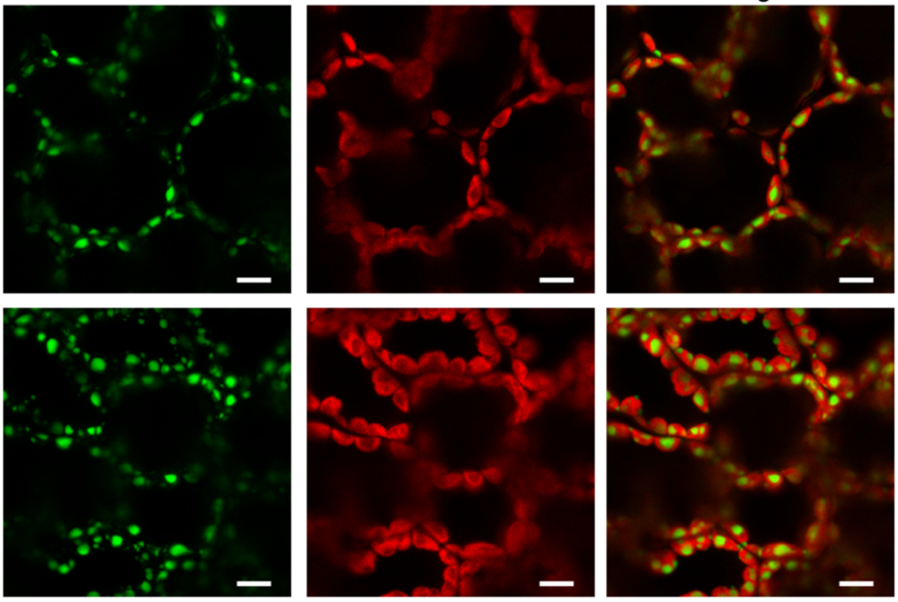


**Figure S2. sfGFP-triggered formation of Rubisco condensates remained unaltered when exposing to dark (top) or 1% $CO_2$ (bottom).** Scale bar: 10 μm

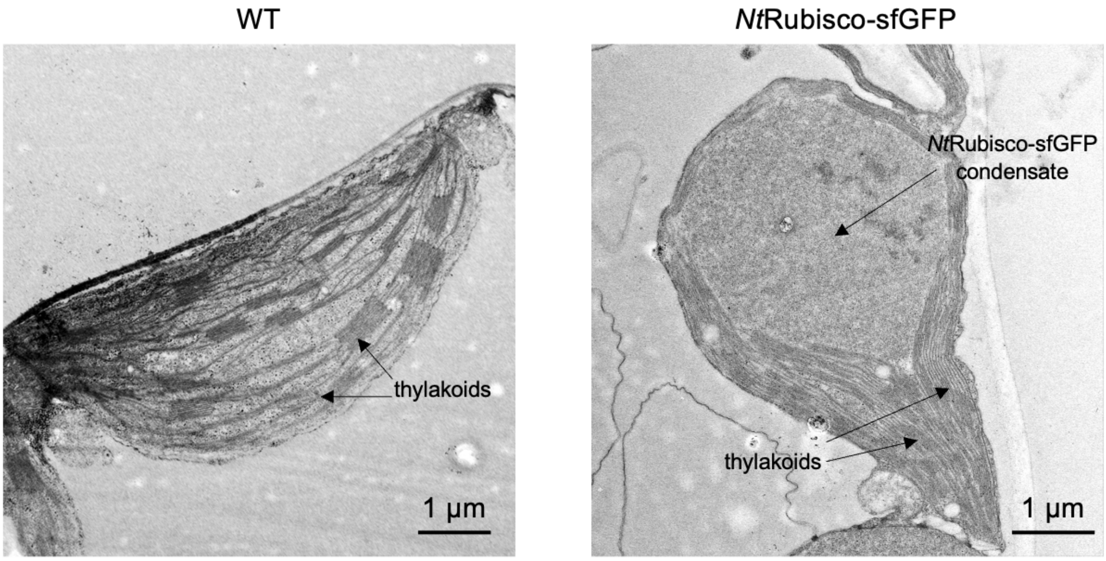


**Figure S3. Transmission electron micrographs of leaf sections of WT and *Nt*Rubisco-sfGFP plants.**

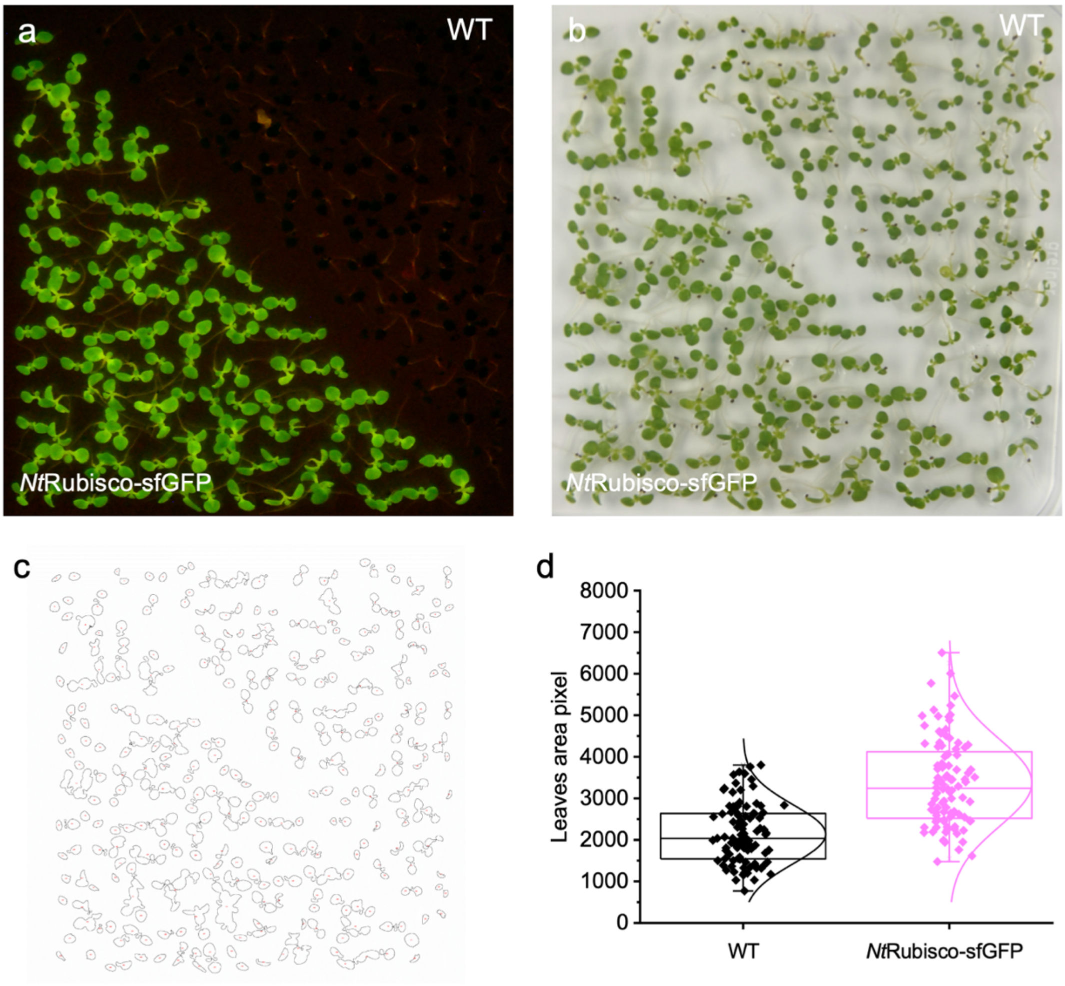


**Figure S4. *Nt*Rubisco-sfGFP showed a slightly larger cotyledon in early stage (15 days after sowing) on MS medium. a** and **b**, Plant images of *Nt*Rubisco-sfGFP and WT, which were taken under blue light (a) and white light (b). **c** and **d**, leaves identified from b by Image J (c) to calculate the cotyledon area (d).

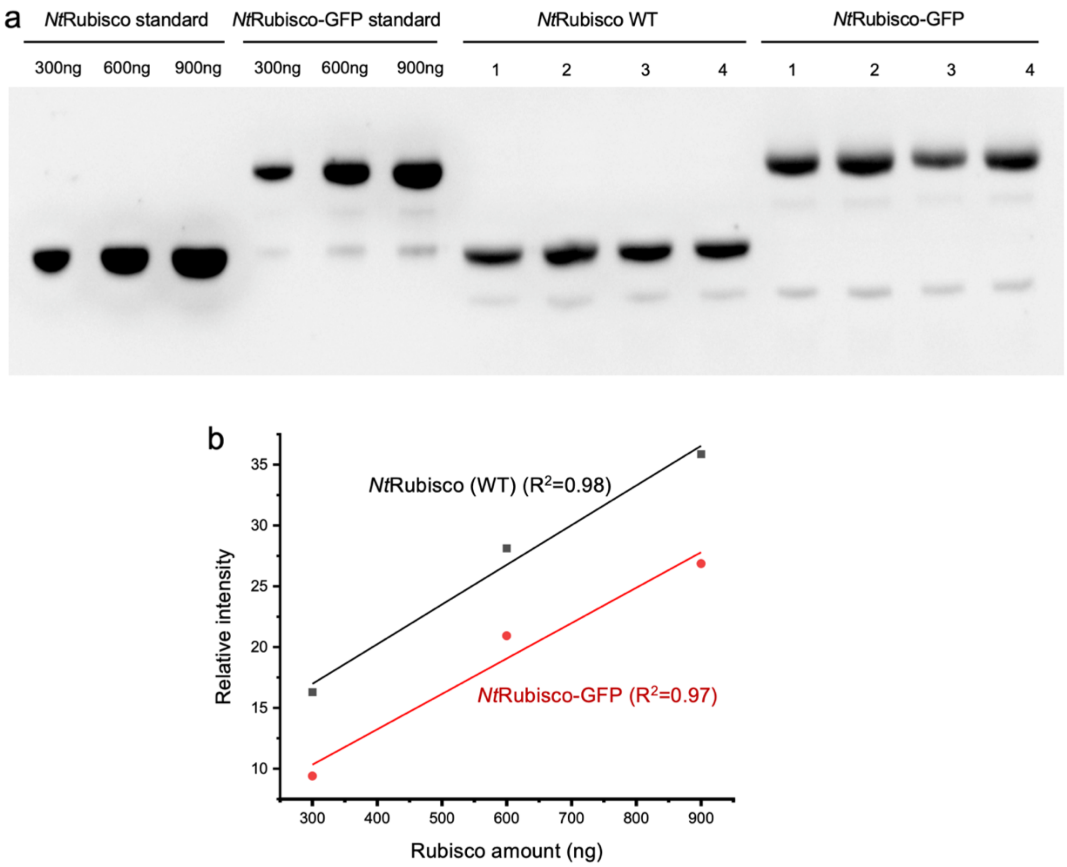


**Figure S5. Quantification of Rubisco content in tobacco leaves by immunoblot analysis using an α-RbcL antibody.** The purified Rubisco from WT and *Nt*Rubisco-sfGFP were quantified by Bradford method and loaded at different concentrations (a) to generate the standard curves (b).
